## Supplementary material for "Repulsive Sema3E-Plexin-D1 signaling coordinates both axonal extension and steering via activating an autoregulatory factor, Mtss1": Key resource table.pdf

### Supplementary Table1

#### KEY RESOURCE TABLE

| REAGENT or RESOURCE | SOURCE | IDNETIFIER |
|---|---|---|
| **Antibodies** | | |
| Rabbit anti-Mtss1 | Novus Biologicals | Cat# NBP2-24716<br>RRID: AB_2716709 |
| Goat anti-Plexin-D1 | R&D systems | Cat# AF4160<br>RRID: AB_2237261 |
| Goat anti-Tau | Santa Cruz | Cat# sc-1995<br>RRID: AB_632467 |
| Mouse anti-Neurofilament | Hybridoma Bank | Cat# 2H3<br>RRID: AB_ 531793 |
| Rabbit anti-β-actin/HRP | Cell Signaling Technology | Cat# 5125S<br>RRID: AB_1903890 |
| Mouse anti-Myc | Cell Signaling Technology | Cat# 2276<br>RRID: AB_331783 |
| Goat anti-Vsv | Abcam | Cat# ab3861<br>RRID: AB_304118 |
| Human anti-Sema3E | LSBio | Cat# LS-c353198 |
| Rabbit anti-Phospho-Akt | Cell Signaling Technology | Cat# 9271<br>RRID: AB_329825 |
| Rabbit anti- Akt | Cell Signaling Technology | Cat# 9272<br>RRID: AB_ 329827 |
| Rabbit anti-RFP | Abcam | Cat# ab62341<br>RRID: AB_945213 |
| Mouse anti-RFP | Thermo Fisher Scientific | Cat# MA5-15257<br>RRID: AB_10999796 |
| Mouse anti-alpha-tubulin | Sigma-Aldrich | Cat# T5168<br>RRID: AB_477579 |

| | | |
|---|---|---|
| Rabbit anti-cleaved caspase3 | Cell Signaling Technology | Cat# 9661<br>RRID: AB_2341188 |
| CD31 | BD Bioscience | Cat# 553370<br>RRID: AB_394816 |
| anti-digoxigenin-alkaline phosphatase | Roche | Cat# 11093274910<br>RRID: AB_2313640 |
| Goat anti-mouse IgG/HRP | Thermo Fisher Scientific | Cat# 31430<br>RRID: AB_228307 |
| Donkey anti-rabbit IgG/HRP | Jackson ImmunoReasearch | Cat# 711-035-152<br>RRID: AB_10015282 |
| Donkey anti-goat IgG/HRP | Jackson ImmunoReasearch | Cat# 705-035-147<br>RRID: AB_2313587 |
| Donkey anti-rabbit IgG, Alexa Fluor 488 | Thermo Fisher Scientific | Cat# A-21206<br>RRID: AB_2535792 |
| Donkey anti-mouse IgG, Alexa Fluor 488 | Thermo Fisher Scientific | Cat# A-21202<br>RRID: AB_141607 |
| Donkey anti-goat IgG, Alexa Fluor 568 | Thermo Fisher Scientific | Cat# A-11057<br>RRID: AB_142581 |
| Donkey anti-rabbit IgG, Alexa Fluor 568 | Thermo Fisher Scientific | Cat# A-10042<br>RRID: AB_2534017 |
| Donkey anti-mouse IgG, Alexa Fluor 568 | Thermo Fisher Scientific | Cat# A-10037<br>RRID: AB_2757558 |
| Donkey anti-mouse IgG, Alexa Fluor 647 | Thermo Fisher Scientific | Cat# A-31571<br>RRID: AB_162542 |
| Chemicals, peptides, and kits | | |
| MK2206 | SelleckChem | Cat# S1078 |
| TRIzol$^{TM}$ Reagent | Thermo Fisher Scientific | Cat# 15596026 |
| QuantiTect Reverse Transcription kit | Qiagen | Cat# 205313 |
| LightCycler 480 SYBR Green I Master | Roche | Cat# 04 887 352 001 |
| RNasin® Ribonuclease Inhibitor | Promega | Cat# N2115 |

| | | |
|---|---|---|
| Halt™ Protease and Phosphatase Inhibitor Cocktail | Thermo Fisher Scientific | Cat# 78444 |
| Pierce™ BCA Protein Assay Kit | Thermo Fisher Scientific | Cat# 23225 |
| Immobilon-P PVDF Membrane | Merck | Cat# IPVH00010 |
| SuperSignal™ West Pico PLUS Chemiluminescent Substrate | Thermo Fisher Scientific | Cat# 34580 |
| SuperSignal™ West Femto Maximum Sensitivity Substrate | Thermo Fisher Scientific | Cat# 34096 |
| ProLong™ Diamond Antifade Mountant with DAPI | Thermo Fisher Scientific | Cat# P36962 |
| Eukitt® Quick-hardening mounting medium | Sigma-Aldrich | Cat# 03989 |
| Alexa Fluor™ 488 Phalloidin | Thermo Fisher Scientific | Cat# A12379 |
| Alexa Fluor™ 568 Phalloidin | Thermo Fisher Scientific | Cat# A12380 |
| Alexa Fluor™ 647 Phalloidin | Thermo Fisher Scientific | Cat# A22287 |
| Gibco™ DMEM, high glucose, pyruvate | Gibco™ | Cat# 11995-065 |
| Fetal Bovine Serum | Hyclone | Cat# SH30084.03 |
| Penicillin-Streptomycin | HyClone | Cat# SV30010 |
| Opti-MEM™ I Reduced Serum Medium | Gibco™ | Cat# 31985070 |
| HyClone Dulbecco's Phosphate Buffered Saline | HyClone | Cat# SH30028.02 |
| HBSS, no calcium, no magnesium | Gibco™ | Cat# 14170120 |
| Paraformaldehyde | Electron Microscopy Sciences | Cat#19202 |
| Magnesium chloride | Sigma-Aldrich | Cat# M8266 |
| Poly-D-lysine hydrobromide | Sigma-Aldrich | Cat# P6407 |
| Corning® Laminin | Corning | Cat# 354232 |
| Lipofectamine™ 2000 Transfection Reagent | Thermo Fisher Scientific | Cat# 11668019 |
| Basic NucleofectorTM Kit | LONZA | Cat# VAPI-1003 |
| NBT/BCIP Ready-to-Use Tablets | Roche | Cat# 11697471001 |
| DiI (1.1-dioctadecyl-3,3,3,3-tetramethyl-indocarbocyanine perchlorate) | Sigma-Aldrich | Cat# 468495 |
| FD Rapid GolgiStain™ Kit | FD neurotechnologies Inc. | Cat# PK401A |

| | | |
|---|---|---|
| Deposited data | | |
| RNA-seq (P5 mice, striatum) | This paper | GEO: GSE196558 |
| Experimental models: Organisms/strains | | |
| Mouse: C57BL/6J | The Jackson Laboratory | Stock# 000664 |
| Mouse: Nestin-Cre | The Jackson Laboratory | Stock# 003771 |
| Mouse: Tie2-Cre | The Jackson Laboratory | Stock# 008863 |
| Mouse: Drd1a-tdTomato | The Jackson Laboratory | Stock# 016204 |
| Mouse: *Mtss1*$^{flox/+}$ | Center for Animal Resources and Development Database (CARD) under permission of Dr. Mineko Kengaku | Card ID#2760 |
| Mouse: *Plxnd1*$^{flox/flox}$ | Obtained from Dr. Chenghua Gu | Kim et al., 2011 |
| Mouse: *Sema3e*$^{+/-}$ | Obtained from Dr. Chenghua Gu | Chauvet et al., 2007 |
| Software and algorithms | | |
| Image J | NIH | https://imagej.nih.gov/ij/ |
| Prism 9 | GraphPad | https://www.graphpad.com/scientific-software/prism/ |
| Image Lab (v5.2.1) | BIO-RAD | https://www.bio-rad.com/ |
| Fusion FX | Vilber | https://www.vilber.com/fusion-fx/ |
| LightCycler®480 (v1.5.1) | Roche | https://lifescience.roche.com/ |
| Leica Application Suite X | Leica | https://www.leica-microsystems.com/ |
| NIS-Elements AR (v4.51.00) | Nickon | https://www.microscope.healthcare.nikon.com/ |
| NIS-Elements (v4.50.00) | Nickon | https://www.microscope.healthcare.nikon.com/ |
| AIVIA | Aivia, Inc. | https://www.aivia-software.com/ |
